# TREM2 macrophages are associated with enhanced response to PD-1 blockade in human hepatocellular carcinoma

**DOI:** 10.1101/2025.09.09.675071

**Authors:** Pauline Hamon, Matthew D. Park, Jessica Le Berichel, Merav Cohen, Brian Y. Soong, Mark Buckup, Clotilde Hennequin, Katherine E. Lindblad, Raphaël Mattiuz, Igor Figueiredo, Alexandra Tabachnikova, Travis Dawson, Darwin D’souza, Leanna Troncoso, Giorgio Ioannou, Colles Price, Nicolas Fernandez, Amir Giladi, Oren Barboy, Zhen Zhao, Sinem Ozbey, Sarah Cappuyns, Amanda Reid, Steven Hamel, Joel Kim, Romain Donne, Christie Chang, Robert Marvin, Hiyab Stefanos, Grace Chung, Raphaël Merand, Laszlo Halasz, Samarth Hegde, Lou M. Guerin, Min Ni, Yi Wei, Gurinder Atwal, Alona Lansky, Hajra Jamal, Nancy Yi, Theodore Chin, Nicola James, Nausicaa Malissen, Fiona Desland, Yonit Lavin, Stephen C. Ward, Maria Isabel Fiel, Rachel Brody, Jeroen Dekervel, Diether Lambrechts, Ephraim Kenigsberg, Edgar Gonzalez-Kozlova, Vladimir Roudko, Alice O. Kamphorst, Jiang He, Marco Colonna, Seunghee Kim-Schulze, Sacha Gnjatic, John C. Lin, Gavin Thurston, Amaia Lujambio, Myron Schwartz, Ido Amit, Thomas U. Marron, Miriam Merad

## Abstract

Macrophages are known to dampen tumor immunity. However, identifying druggable targets that modulate these cells to improve existing immunotherapies has been limited by a dearth of studies identifying macrophages that associate with pathological response to immune checkpoint blockade. To fulfill this unmet clinical need, we leveraged transcriptional and spatial profiling of specimens collected from a Phase II clinical trial studying neoadjuvant PD-1 blockade in patients with hepatocellular carcinoma (HCC). We determined that the intratumoral abundance of TREM2-expressing macrophages and serological levels of soluble TREM2 are elevated in patients who responded to PD-1 blockade, compared to non-responders. We validated these findings in a second HCC cohort and in the IMbrave150 trial. These highlight the robust potential for TREM2 macrophages to predict therapeutic responses of HCC to immunotherapy. Therefore, our study provides a novel basis for the use of TREM2 macrophages to strategize treatment for patients with HCC to maximize therapeutic benefit.

## INTRODUCTION

Given its dismal survival rate and the paucity of treatment options, hepatocellular carcinoma (HCC) constitutes a major global health concern ^1,2^. Surgical resection or ablation remain the standards of care for patients with early-stage disease; however, the majority of patients experience a recurrence of their disease, predominantly in the liver ^3^. This is commonly attributed to the persistence of residual tumor cells that re-grow following surgery. Therefore, intervening with anti-tumoral biologics, such as immune checkpoint blockade ^4–7^, prior to surgery may enhance the curative efficacy of removing resectable tumors ^8–10^. Meeting this objective requires the identification of cellular determinants of response to immune checkpoint blockade to optimize the combination of biologic therapies for neoadjuvant use.

In a phase II trial (NCT03916627), in which patients with early-stage, treatment-naïve HCC received two doses of cemiplimab prior to surgical resection, 30% of patients exhibited pathologic response (complete and partial) ^11^. We found that the tumor microenvironments (TME) of these responsive patients (responders) included both T cell-enriched and -deprived landscapes. Importantly, a subset of non-responders also exhibited T cell-enriched TME. Therefore, we sought to identify lymphoid determinants of response to PD-1 blockade that distinguished T cell-enriched responders and non-responders and determined that a topographical congregation of CXCL13^+^ helper T cells and PD-1^HI^ effector CD8 T cells is uniquely expanded in the tissues of responders^12^.

We recently demonstrated that upon exposure to tumor debris, macs upregulate the Triggering Receptor Expressed on Myeloid Cells 2 (TREM2), which drives the expression of many co-regulated genes (*Gpnmb, Apoe, Cd9, Fabp5, Lipa*) that reduced antitumor immunity by limiting NK cell infiltration and effector function in a non-small cell lung cancer (NSCLC) model ^13^. Similar TREM2 macs were shown to limit antitumor T cell immunity in pre-clinical models ^14–18^. Paradoxically, TREM2 macs demonstrated protective roles in metabolic disorders, liver pathologies, and Alzheimer’s disease ^19–23^, raising essential questions about their role in HCC pathogenesis and response to PD-1 blockade, and whether TREM2 blockade would be beneficial or harmful in these patients.

In the present study, we characterized the myeloid compartment of these patients to identify the cell types that likely regulate T cell activity during PD-1 blockade. Leveraging both single-cell mRNA and spatial transcriptomics, we identified a novel role for TREM2-expressing macrophages in the positive response to PD-1 blockade. Across multiple cohorts of patients, we also demonstrate the utility of soluble TREM2 as a biomarker of immunotherapy response in patients with HCC. Together, these results define a novel myeloid-based stratagem for exploiting the neoadjuvant therapeutic window for immune checkpoint blockade.

## RESULTS

### Distinct monocyte-derived macrophage programs are specifically enriched in HCC lesions that respond to PD-1 blockade

To identify immunological correlates of response to PD-1 blockade, we analyzed 36 surgical resections; these included those from both treatment-naïve patients and patients who received two doses of cemiplimab ^11^ or two to ten doses of nivolumab (off the clinical trial) (**Fig. 1a**). Pathological response was determined for each tissue specimen, with complete response defined as ≥70% tumor necrosis and partial response as ≥50% tumor necrosis. We performed single-cell mRNA sequencing (scRNAseq) of tumor lesions and adjacent, tumor-free tissues to create an atlas of 786,832 immune cells. Of these, using orthogonal iterations of unsupervised clustering ^12,24^, we identified three broad sublineages of mononuclear phagocytes (MNPs), including macrophages (36,616 cells), dendritic cells (20,674 cells), and monocytes (29,528 cells) (**Fig. S1a-b**). Collectively, macrophages were uniquely enriched in tumors, compared to the tumor-free, adjacent tissue (**Fig. S1c**). And among these phagocytes, we found that the tissue-resident Kupffer cells and macrophages derived from adult bone marrow monopoiesis (monocyte-derived macrophages, mo-macs) were present at comparable frequencies in tissues from treatment-naïve, responsive, and non-responsive patients (**Fig. 1b**). Among the monocyte populations, we identified CD14 classical monocytes (*CD14, S100A12, THBS1*), intermediate monocytes, and CD16 non-classical monocytes (*FCGR3A, LYPD2, CDKN1C*) (**Fig. S1d**). Notably, we also detected a distinct subset of tissue-infiltrating monocytes expressing *SPP1, CCL2, FN1,* and *IL4I1*, which were significantly enriched in tumor lesions of responders (**Fig. S1e-f**). Unlike circulating monocytes in peripheral blood, these tumor-infiltrating CD14 monocytes exhibited a pro-inflammatory transcriptional profile, with high expression of *NLRP3, IL1B, CCL3, CCL4,* and *CXCL8* (**Fig. S1g-h**).

**Figure 1:**
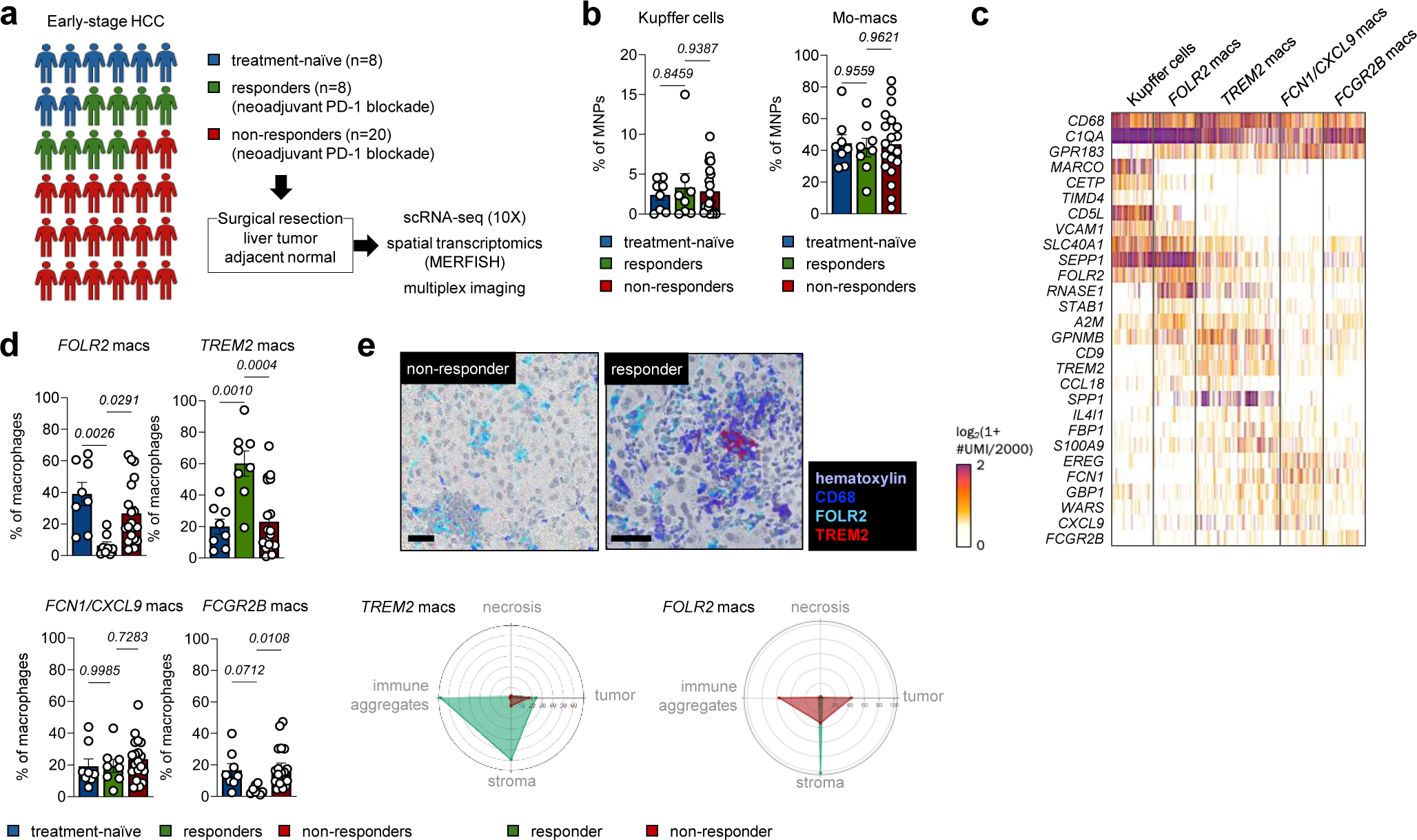
Distinct monocyte-derived macrophage programs are specifically enriched in HCC lesions that respond to PD-1 blockade. **a,** Schematic of the cohort of patients recruited to study early-stage HCC in treatment-naïve patients and after receiving neoadjuvant anti-PD-1 antibody. **b,** Histogram of percentage of Kupffer cells (left panel) and mo-macs (right panel) macs in treatment-naïve, responders and non-responders to PD-1 blockade in tumor lesions. Dots represent percentage of MNPs per tumor samples ± SEM. One-way ANOVA followed by Dunnett’s multiple comparison test was performed. **c,** Heatmap representing single-cell gene expression in mac clusters from both tumor and adjacent tissues. **d,** Histogram of percentage of macs in treatment-naïve, responders and non-responders to PD-1 blockade in tumor lesions for each subset of macs. Dots represent percentage of macrophages ± SEM. One-way ANOVA followed by Dunnett’s multiple comparison test was performed for each subset. **e,** Representative images of multiplex IHC of tumor lesions stained for macs (CD68, FOLR2, and TREM2) and counterstained with hematoxylin. Scale bars represent 50µm. Quantification of TREM2 mac (CD68+ TREM2+), and FOLR2 macs (CD68+ FOLR2+) abundance using multiplex imaging in the different tissue area in responders and non-responders, including tumor nodules, stroma, necrotic areas and immune aggregates.

Given the preponderance of macs in tumor tissues of patients, regardless of treatment or response status, we sought to shed greater light on the complex heterogeneity of mo-macs as it may reveal cell states that are individually associated with response. Altogether, we identified four major transcriptional states, apart from the cells bearing the resident Kupffer cell (KC) mRNA signature (i.e., *MARCO, CD5L, TIMD4, LYVE1, VCAM1, CETP, IFI27, CFP*). These other four broadly expressed mo-mac genes (i.e., *CD68, C1QA, C1QB, C1QC, CD163, GPR183*): (1) FOLR2 macs (i.e., *FOLR2, SEPP1, SLC40A1, F13A1, STAB1, IGF1, GPR34, RNASE1*), (2) TREM2 macs (i.e., *TREM2, GPNMB, SPP1, NUPR1, APOE, FABP4, FABP5, CAPG, CD9*), (3) FCN1/CXCL9 macs (i.e., *FCN1, THBS1, IL1R1, GPB1, PLAUR, CCL20, CXCL9, CXCL10*), and (4) FCGR2B macs (i.e., *CLEC10A, CD1C, CD1E*) (**Fig. 1c**, **Fig. S1i**, **Fig. S2a-b**).

As previously described for tissue-resident alveolar macrophages (AMs) in non-small cell lung cancer lesions (NSCLC) ^24–26^, KCs were significantly reduced in tumors, compared to adjacent, tumor-free tissue (**Fig. S1j**). In contrast, FOLR2 and TREM2 macs were all enriched in tumor lesions (**Fig. S1j**); FCN1/CXCL9 macs were present at comparable frequencies in tumoral and adjacent, normal tissue, whereas FCGR2B macs were preferentially enriched in the normal tissue. Notably, FOLR2 macs represented the most abundant subset in the tumors of treatment-naïve and non-responsive patients and was substantially under-represented in tumors of responders. However, TREM2 macs were significantly enriched in tumors of responders, compared to either treatment-naïve patients or non-responders (**Fig. 1d**, **Fig. S2c**). Given their enrichment in tumor tissues (albeit differently by response status), we sought to refine our study of the FOLR2 and TREM2 macs as potential regulators of response.

We next sought to characterize the spatial organization of these macs within the TMEs of the treatment-naïve, responsive, and non-responsive patients. We first annotated tissue sections histologically, defining areas of tumor, stroma, necrosis, and immune aggregation (**Fig. S2d**). Comparing the tissue topography across the patient groups based on these criteria revealed that high stromal and necrotic tissue content at the time of surgical resection is associated with response to PD-1 blockade (**Fig. S2e**). Then, immunohistochemical staining of TREM2 and FOLR2 enabled protein-level validation and spatial mapping of these mac subsets within the tumor tissue. We confirmed that TREM2 macs were enriched in responders, compared to non-responders, and observed that their specific accumulation within the intratumoral immune aggregates and the stroma (**Fig. 1e**). In comparison, FOLR2 macs were found to be sequestered in the stroma of responders, whereas in non-responders, FOLR2 macs were more enriched in the immune aggregates and tumor-occupied areas. Given the specificity of *TREM2* mRNA to the mo-mac sublineage (**Fig. S2a-b**) and the associations of TREM2 macs with response, we hypothesized that these cell states may help orchestrate response in patients HCC receiving PD-1 blockade.

### TREM2 macs are uniquely enriched in proximity to PD-1^hi^ CD4 helper T cells

To determine how these macs may help determine response to PD-1 blockade, we applied the PIC-seq platform (physically-interacting cell sequencing) to identify the cell types that interact directly with macs in tumors and in adjacent normal tissues ^27,28^. We directed our focus to examining the T cells that may be involved in these interactions. We captured a greater number of these MNP/T cell PICs in tumors, compared to normal tissues (**Fig. 2a-b**). By annotating the sequenced singlet MNPs, we deconvolved the MNPs sequenced amongst the PICs. Strikingly, only TREM2 macs were found to be enriched in PICs compared to singlets (**Fig. 2a-b**), highlighting a unique role that they may play in directly modulating tumor-infiltrating T cells and their response to PD-1 blockade.

**Figure 2:**
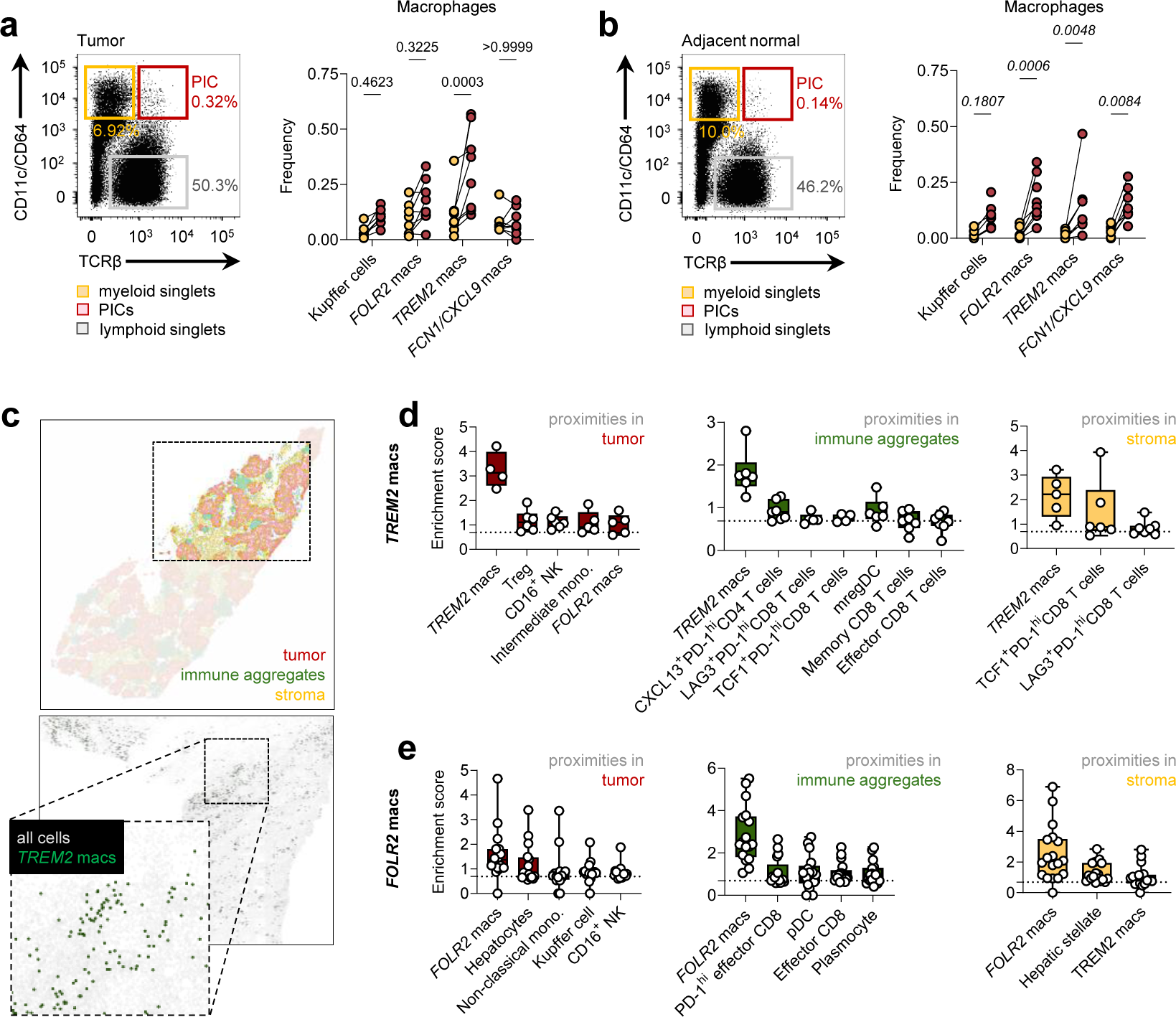
TREM2 macs are uniquely enriched in proximity to PD-1^hi^ CD4 helper T cells. **a,** Representative dot plot of the singlets of T cells and MNP and the PIC populations in tumor lesion. Quantification of MNP singlet (yellow) and PIC (red) for each of the mac subsets in tumor lesions. Dots represent mean ± SEM per patient. Two-way ANOVA followed by Sidack multiple comparison was performed. **b,** Representative dot plot of the singlets of T cells and MNP and the PIC populations in adjacent tissue. Quantification of MNP singlet (yellow) and PIC (red) for each of the mac subsets in adjacent tissues. Dots represent mean ± SEM per patient. Two-way ANOVA followed by Sidack multiple comparison was performed. **c,** Representative image of computational annotation of MERFISH responder sample to define tumor, stroma and immune aggregate area. Computational projection and spatial identification of *TREM2* macs on MERFISH HCC slide of a responder. **d,** Proximity analysis of immune subsets significantly enriched in a ≤30µm distance of TREM2 macs in defined area including tumor nodules (left panel), immune aggregates (middle panel) and stroma (right panel). **e,** Proximity analysis of immune subsets significantly enriched in a ≤30µm distance of FOLR2 macs in defined area including tumor nodules (left panel), immune aggregates (middle panel) and stroma (right panel).

Limited by our ability to deconvolve the T cells in PICs, we applied an orthogonal spatial transcriptomics-based approach. The multiplexed error-robust fluorescence *in situ* hybridization (MERFISH) technology combines single-cell mRNA detection with higher spatial resolution. Using the gene signatures defined from our scRNAseq dataset, we identified mo-mac subsets (**Fig. S3a**), and defined tumor regions based on expression of tumor cell and hepatocyte genes, immune aggregates based on the expression of B cell genes, and stroma based on the expression of fibroblast and hepatic stellate cell (HSC) genes (**Fig. 2c**, **Fig. S3b**). Combining these annotations with the identification of MNPs based on the single-cell mRNA detection from the MERFISH technology helped maximize our insight into both the spatial organization and intercellular interactions between TREM2 macs and FOLR2 macs with T cells within the TME. In tumor nodules, TREM2 macs were most proximal to Tregs and NK cells, as well as monocytes and FOLR2 macs (**Fig. 2d**, left); whereas FOLR2 macs were primarily associated with non-classical monocytes and NK cells (**Fig. 2e**, left). Within immune aggregates, FOLR2 macs were found in close proximity to PD-1^hi^ Effector CD8 T cells and Effector CD8 T cells (**Fig. 2e**, middle). In contrast, TREM2 macs were spatially organized near multiple subsets of PD-1^hi^ T cells, including CXCL13^+^ PD-1^hi^ CD4 T cells, TCF1^+^ PD-1^hi^ CD8 T cells, and LAG3^+^ PD-1^hi^ CD8 T cells (**Fig. 2d**, middle, **Fig. S3c**). Consistent with our prior study, where these T cells were shown to associate with mature DCs and response to PD-1 blockade ^12^, mature DCs (mregDCs) were also found proximal to TREM2 macs in immune aggregates. In the stroma, TREM2 macs were also found in close proximity to TCF1^+^ PD-1^hi^ CD8 T cells and LAG3^+^ PD-1^hi^ CD8 T cells (**Fig. 2d**, right). Meanwhile, FOLR2 macs were found close to hepatic stellate cells and TREM2 macs (**Fig. 2e**, right).

To further explore the functional implications of these spatial interactions, we examined transcriptional differences in TREM2 macs and PD-1^hi^ CD8 T cells based on their proximity to one another. Differential gene expression analysis revealed that PD-1^hi^ CD8 T cells in direct contact with TREM2 macs exhibited elevated levels of activation and cytotoxicity markers (*KLRK1, CD69, CD27, CXCR6, CXCR4, CD2*), as well as the transcription factor *EOMES* and cytotoxic cytokines (*CXCL16, GZMA, GZMK, GZMH*) (**Fig. S3d**). In contrast, those distal to TREM2 macs showed increased expression of exhaustion markers (*HAVCR2, TOX, TIGIT*). Similarly, TREM2 macs engaged in direct interactions with PD-1^hi^ CD8 T cells exhibited enhanced antigen-presenting capacity, marked by increased expression of MHC-II molecules (*HLA-DQA1, -DQB1, -DRB1, -DRA*), cytokine receptors, and T cell co-stimulatory molecules (*IFNGR1, CD86*) (**Fig. S3e**). TREM2 macrophages distant from PD-1^hi^ CD8 T cells upregulated markers associated with alternative mac subsets (*MARCO*, defining Kupffer cell identity, and *FCN1*, associated with FCN1 macs), which are not typically linked to immunotherapy response. Altogether, these data show that TREM2 macs may directly influence the anti-tumor immune response and determine sensitivity to PD-1 blockade, given their close interactions with PD-1^hi^ T associated with immune activation and enriched in responders ^12^.

### Regulators of phagocytosis distinguish hepatic TREM2 macs from similar cells in other tumors

TREM2 macs have been described in various tumor types, including NSCLC, breast cancer, and ovarian cancer; in these settings, TREM2 macs have been ascribed a immunosuppressive function that promotes tumor progression ^14,16,24^. Using a preclinical model of lung cancer, we also previously demonstrated that TREM2 macs deter NK cell infiltration and cytotoxic activity and blocking TREM2 can salvage anti-tumor immunity ^13^. In HCC, however, TREM2 macs have been linked to anti-tumoral activity. Given our findings, which suggests a novel protective function of TREM2 macs in the context of immunotherapy response, we leveraged published scRNAseq datasets (advanced HCC, NSCLC, pancreatic cancer, and triple-negative breast cancer) to determine whether the TREM2 program from the present study also describes mo-macs of other solid tumors ^24,29–31^. This analysis revealed that our TREM2 program defines TREM2 macs that heavily populate these other types of tumors (**Fig. 3a**). To further establish that these macs are, in fact, transcriptionally concordant, we computed differentially-expressed genes that distinguish TREM2^hi^ from TREM2^lo^ mo-macs in HCC and in NSCLC. Comparing these lists between the HCC and NSCLC datasets identified 48 conserved genes constituting a core TREM2 program, as well as 209 liver-specific genes and 89 lung-specific genes (**Fig. 3b**). Projecting the conserved gene signature onto mo-macs from the other datasets demonstrated that it indeed defines mo-macs of other tumor types (**Fig. S3f**). Since the TREM2 program is induced in naïve bone marrow-derived macs (BMDMs) when cultured with apoptotic cell debris, our findings suggest that the core TREM2 program is likely induced by cues common to TMEs of different tumor types. In contrast, the opposing roles of TREM2 macs in anti-tumor immunity – and, in this case, response to PD-1 blockade – between HCC and NSCLC, for example, is likely determined by the tissue-specific DEGs.

**Figure 3:**
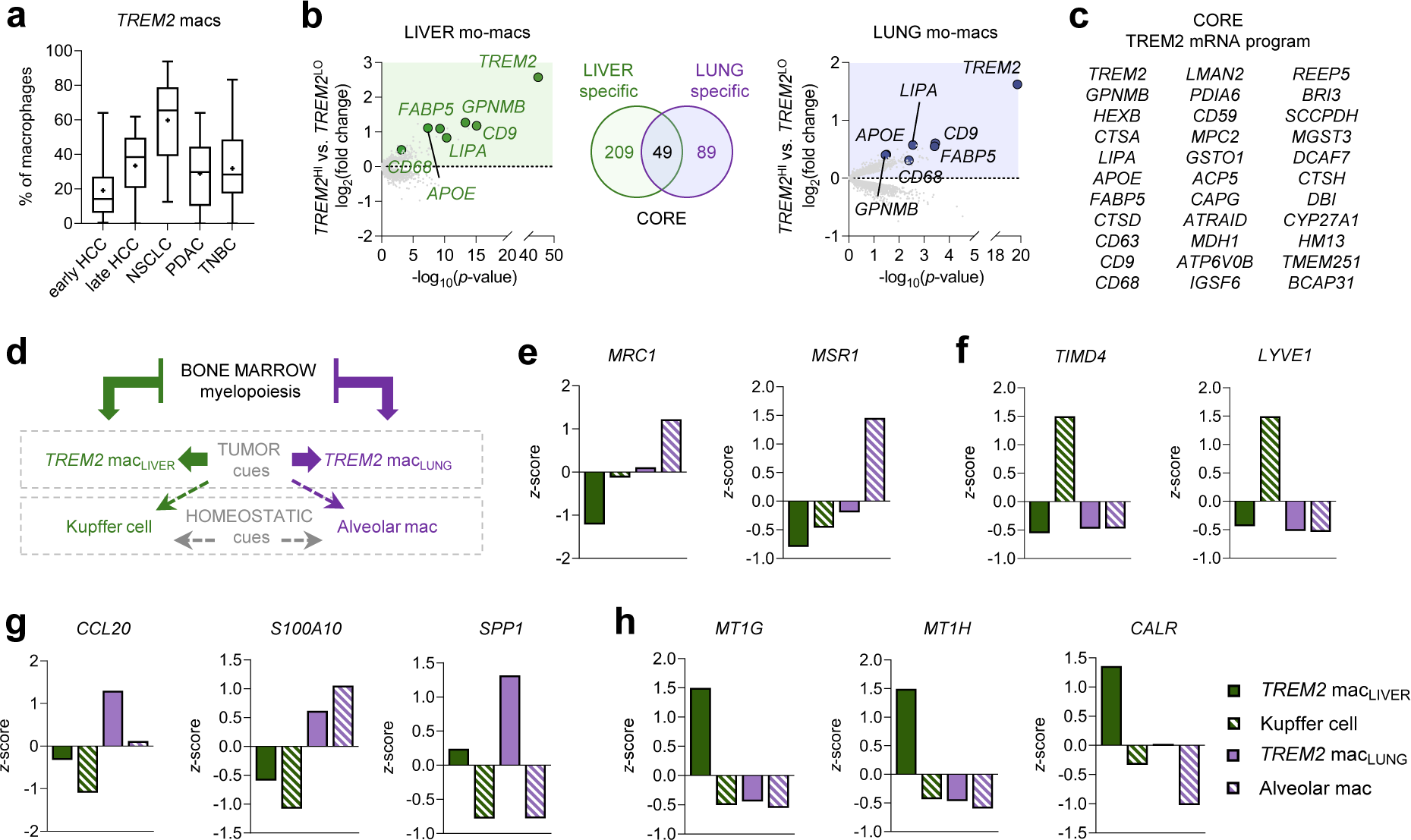
Regulators of phagocytosis distinguish hepatic TREM2 macs from similar cells in other tumors. **a,** Percentage of macs expressing *TREM2* across multiple tumor types (HCC: Hepatocellular carcinoma, NSCLC: Non-small cell lung cancer, PDAC: Pancreatic ductal adenocarcinoma, TNBC: Triple negative breast cancer). **b,** Differentially expressed genes (DEGs) between *TREM2^Hi^* versus *TREM2^Lo^* macs in HCC and NSCLC. **c,** Genes conserved by *TREM2^Hi^* macs in both HCC and NSCLC. **d,** Scheme of the approach to understand the tissue versus tumor cues and the impact of myelopoiesis in the TREM2 mo-mac program versus RTM. **e-h,** Analysis of transcripts expression significantly enriched between TREM2 macs in NSCLC and HCC or the RTM (Alveolar macs in the lung and Kupffer cells in HCC), showing z-score of gene expression either enriched in RTM **(e)**, TREM2 macs in NSCLC **(g)** and TREM2 macs in HCC **(h)**.

To properly contextualize these genes, we further computed tissue-specific genes that not only distinguish the TREM2 macs in HCC from those in NSCLC but also account for the difference between these mo-macs generated during chronic inflammation associated with tumorigenesis and resident tissue macrophages (RTMs) responsible for homeostasis (i.e., AMs in the lungs and KCs in the liver) (**Fig. 3d**). This analysis validated the use of prototypical cell-surface markers of AMs (i.e., *MRC1, MSR1*) and KCs (i.e., *TIMD4*, *LYVE1*) (**Fig. 3e-f**). TREM2 macs in NSCLC expressed higher levels of immunomodulatory chemokines (i.e., CCL20) ^32,33^ and TLR regulators (i.e., S100A10) ^34^, as well as SPP1, a matricellular protein associated with tissue remodeling ^35^ (**Fig. 3g**). In contrast, hepatic TREM2 macs expressed highest levels of metallothionens (i.e., *MT1G, MT1H*), which are essential for protecting against oxidative stress and inhibiting ferroptosis-related lipid peroxidation ^36–39^ (**Fig. 3h**). Notably, TREM2 macs in HCC also expressed uniquely high levels of *CALR*, which encodes calreticulin (**Fig. 3h**). This molecule on the cell surface has been to facilitate clearance of cell debris, counteracting the effects of CD47 and its anti-phagocytic function ^40–42^. These unique features suggest that TREM2 macs in the liver are equipped to dampen pro-tumorigenic cytotoxic stress signals and effectively remove inflammatory cell debris. In sum, these key transcriptional differences reflect the tissue- and tumor-specific imprints that may help TREM2 macs specialize in distinct immunological functions in different tumor types.

### The TREM2 program predicts response to ICB and overall survival of patients with HCC

To validate that the TREM2 program associates with response to PD-1 blockade, we analyzed an independent cohort of patients treated with cemiplimab and stereotactic body radiotherapy (SBRT) in the neoadjuvant setting. Of the 21 patients recruited to this cohort, 6 patients responded to treatment. We performed scRNAseq on tumor and adjacent normal tissue from 16 patients. Consistent with our findings from the exploratory cohort, we observed an enrichment of mo-macs and an exclusion of KCs from tumor samples. Among mo-macs, TREM2 macs were specifically enriched in responders (**Fig. 4a**). Importantly, across both the exploratory and validation cohorts, responders, including partial responders, exhibited a lower recurrence rate, compared to non-responders, which may suggest that TREM2 macs play a critical role in prolonging relapse-free survival (**Fig. 4b**). Finally, to further enhance the statistical strength of our work, we analyzed data from the IMbrave150 phase III trial, in which patients with HCC (n=358) were treated with the combination of atezolizumab (anti-PD-L1) and bevacizumab (anti-VEGF) ^43^. We used the top 10 genes of the TREM2 mRNA program to generate a transcriptional score from the bulk RNA-seq of patient tumor samples, then stratify patients into high (TREM2^hi^; top quartile) and low (TREM2^lo^; bottom quartile) scorers. Notably, TREM2^hi^ patients exhibited a significant survival benefit, compared to their TREM2^lo^ counterparts (**Fig. 4c**, top). To assess whether this survival advantage extended to other cancer types, we applied the same method to the bulk RNA-seq data generated from the POPLAR phase II trial, in which patients with lung cancer were treated with atezolizumab ^44^. However, unlike our observations from the IMbrave150 trial, the TREM2 program failed to stratify patients with significant differences in response to PD-L1 blockade and overall survival (**Fig. 4c**, bottom), reinforcing the principle that the immunological function of TREM2 macs is tissue-dependent.

**Figure 4:**
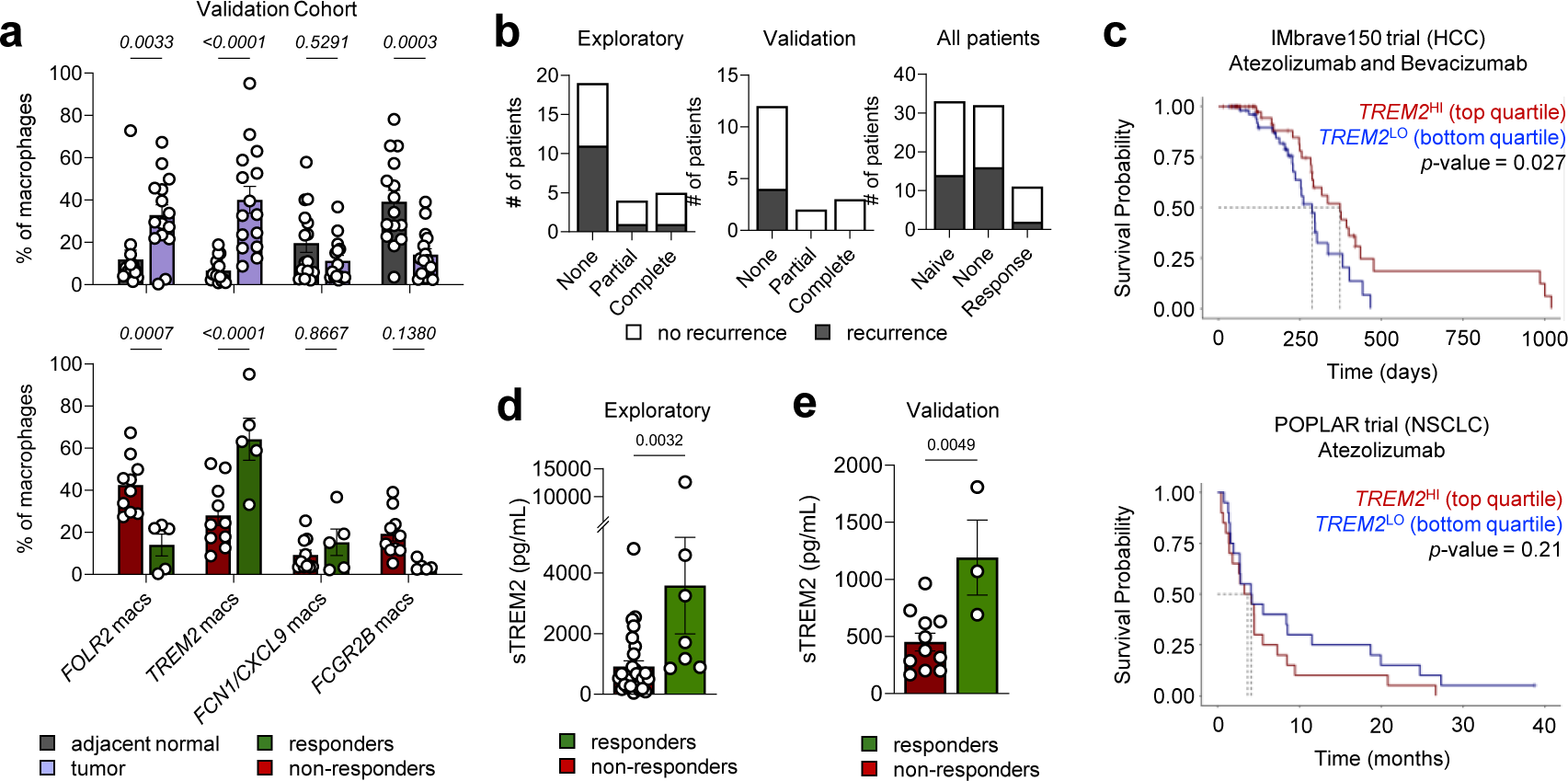
The TREM2 program predicts response to ICB and overall survival of patients with HCC. **a,** Histograms of percentage of macs in tumor and adjacent tissue for each subset of mo-macs. Dots represent percentage of macrophages ± SEM. Two-way ANOVA followed by Sidack multiple comparison was performed. **b,** Histograms represent the recurrence of patients in the intial cohort of 36 patients, in the validation cohort and for all the patients. **c,** Kaplan-Meyer survival of top and bottom quartile of patients based on the top 10 genes of the TREM2 signature, in the IMbrave150 and POPLAR trials. **d-e,** Concentration of soluble TREM2 at baseline in responders and non-responders of the exploratory **(d)** and validation **(e)** cohorts. Dots represent sTREM2 concentration ± SEM. Unpaired t-test was performed for each graph.

Finally, our findings reveal a striking association between TREM2 macrophages and response to PD-1 blockade. By quantifying soluble TREM2 (sTREM2) in baseline serum samples before treatment, we discovered that responders exhibited significantly higher sTREM2 levels compared to non-responders (**Fig. 3d, Fig. S4a-b**). Remarkably, this trend was also seen in our independent validation cohort, where patients with elevated baseline sTREM2 responded significantly better to anti-PD-1 therapy combined with SBRT (**Fig. 3e**). These compelling results position baseline sTREM2 as a promising and easily accessible blood-based biomarker, with the potential to transform patient stratification and enhance precision in PD-1 blockade therapies.

## DISCUSSION

In this study, we found that TREM2 macs and their release of soluble TREM2 are cellular and molecular markers of response to PD-1 blockade for patients with HCC. Our findings are supported by previous studies describing the anti-tumorigenic, protective function of TREM2 macs in the progression of cirrhosis and hepatic cancer ^20,45–48^. The present study, however, presents the first evidence to-date of myeloid TREM2 as a major determinant of response to ICB in HCC. The clinical importance of this study is further highlighted by the fact that TREM2 macs execute different functions in the lungs ^13^ and in the brain ^49–51^, despite their proposed shared ontogeny (i.e., adult bone marrow monopoiesis) ^13,38,52^.

TREM2 is a transmembrane receptor on the cell surface that binds a diverse array of ligands, including lipo-proteins and phospholipids, typically exposed on the surface of apoptotic cell blebs or released cell debris. The uptake of these cargo has been shown to induce the TREM2 mRNA program in professional phagocytes ^13^, including tissue-infiltrating mo-macs. This may likely explain why similar TREM2 programs can be identified from single-cell characterization of injured tissues. We hypothesize that tissue-specific signals – like homeostatic cues derived from hepatocytes versus alveolar epithelial cells – can program an added layer of molecular changes in TREM2 macs that underscore their distinct immunological properties in the liver versus the lungs. The degree of chronic inflammation, such as that seen in HCC, could further influence the activation and programming of TREM2 macrophages, potentially modifying their pro-tumorigenic or immune-modulatory properties. For instance, in the liver, chronic inflammation associated with NASH or fibrosis may induce a different macrophage response. Furthermore, the ferroptosis pathway, which has been shown to protect liver tissue against oxidative stress and lipid peroxidation, may confer protective effects in the liver but not in the lung. Further work must be done to identify the cues responsible for these differences that result in either protective or pathogenic outcomes, which have been demonstrated in pre-clinical models of HCC. For example, recent studies using a carcinogen-induced model of HCC in *Trem2*^−/−^ mice exhibited increased tumor burden, when compared to tumor development in wild-type mice ^45,46^.

In addition to the TREM2 state, our study, alongside others, has highlighted FOLR2 macrophages as integral components of the HCC microenvironment. We found that these cells accumulate within tumor nodules and are enriched in non-responders to PD-1 blockade. Li et al. previously characterized an ‘onco-fetal’ niche comprised of FOLR2 macs, cancer-associated fibroblasts (CAFs), and a subset of endothelial cells, which was associated with cancer relapse ^38,53^. Notably, the enrichment of FOLR2 macrophages in lesions from relapsed patients aligns with our findings, reinforcing the idea that specific macrophage subsets influence disease progression and treatment outcomes. Interestingly, a similar group of FOLR2 macs was identified in breast cancer, where these cells were instead linked to enhanced T cell infiltration and improved survival^54^. This contrast underscores the context-dependent nature of macrophage programs beyond just the TREM2 program, emphasizing the urgent need to dissect the mechanisms that govern their functional specialization across different tissues and tumor types. Understanding how these programs are shaped by and, in turn, shape their local microenvironment will be critical for designing more effective therapeutic strategies.

The immune mechanisms underlying response to therapy are undoubtedly complex. We previously showed that, among patients with high T cell infiltration – which is typically associated with a better prognosis – only half responded favorably to PD-1 blockade^12^. A deeper analysis of these two groups revealed that response was driven by the presence of specific immune niches. In responders, PD-1^hi^ progenitor (TCF1+) CD8 T cells were reactivated and locally expanded through interactions with CXCL13+ T helper cells and mature dendritic cells (mregDC). This triad was essential for sustaining the effector function of PD-1^hi^ CD8 T cells. In contrast, non-responders with high T cell infiltration lacked this triad and instead harbored a higher proportion of terminally exhausted CD8 T cells. Notably, our current study reveals that TREM2 macs closely surround these immune niches and are major partners in direct interactions with PD-1^hi^ CD8 T cells, too. These findings suggest that they may also be integral members of the triad we previously described, so this quartet should be considered as the defining cellular determinants of immunotherapy response. Further investigation is needed to determine whether TREM2 macs directly contribute to the initial formation and/or stabilization of these immune aggregates and how they influence the balance between effective T cell responses and exhaustion.

Altogether, our study opens several avenues for future research. Further investigation into the precise mechanisms through which TREM2 mac physically modulate PD-1^hi^ T cells and direct their response to PD-1 blockade may require the experimental screening of ligand-receptor pairs. Alternatively, paracrine signaling via cytokines and/or metabolites processed and released by TREM2 macs may also facilitate how direct interactions between TREM2 macs and both CD4 and CD8 T cells determine response. So, enhanced metabolic profiling of the T cells involved in these interactions could be key. These longer-term goals will help build the rationale for determining the degree of pre-treatment infiltration of tumors by TREM2 macs to identify the subsets of patients will benefit most from neoadjuvant therapy with immune checkpoint blockade prior to surgical resection.

## MATERIALS AND METHODS

### Human subjects

Samples of tumor, non-involved adjacent liver and peripheral blood mononuclear cells (PBMC) were obtained from patients undergoing surgical resection at Mount Sinai Hospital (New York, NY) after obtaining informed consent in accordance with a protocol reviewed and approved by the Institutional Review Board at the Icahn School of Medicine at Mount Sinai (IRB Human Subjects Electronic Research Application 18-00407) and in collaboration with the Biorepository and Department of Pathology. Demographic details and information linking samples to patients are provided in Table S1.

A single-arm, open-label, phase II clinical trial (ClinicalTrials.gov identifier: NCT03916627, Cohort B) was conducted in patients with resectable HCC. In this cohort, 20 patients received two cycles of cemiplimab prior to undergoing surgical resection, as previously described in Marron et al ^11^. As a validation cohort, we analyzed an independent group of 21 patients enrolled in a subsequent cohort of the same trial (Cohort B2), who were treated with a neoadjuvant regiment combining cemiplimab and stereotactic body radiotherapy (SBRT); scRNAseq data were available for 16 of these patients. Additionally, we included 9 patients treated off-label with 2 to 4 doses of nivolumab before surgery, and 8 treatment-naïve patients who had not receive any tumor-related treatment prior to surgery. For the sera analysis at baseline, we included samples from 17 patients enrolled in a separate clinical trial (NCT04123379, Arms C, D, E).

Response to treatment was defined as more than 50% (partial response) or more than 70% (complete response) necrosis of resected tumor by pathologists. Post-operatively patients underwent standard-of-care screening every 6 months; recurrence was defined as development of imaging consistent with recurrent HCC using the Liver Imaging Reporting and Data Systems (LIRADS) score of 5 for an intrahepatic lesion (the most common site of HCC recurrence) or arterial enhancing new extrahepatic metastatic foci consistent with metastatic disease.

We analyzed patient data from two external clinical trials: IMBRAVE150 (NCT03434379)^43^ and POPLAR (NCT01903993)^44^. The IMbrave150 trial assessed atezolizumab (anti-PD-L1) plus bevacizumab (anti-VEGF) in patients with unresectable HCC. The POPLAR trial investigated Atezolizumab versus Docetaxel in NSCLC patients.

### Tissue processing

Freshly resected specimens were transported to the lab on ice in RMPI media supplemented with 10% FBS. Each sample was subdivided: a fragment of tissue was formalin-fixed and paraffin embedded (FFPE) for histology, another fragment for frozen for RNA/DNA detection and the remaining tissue was processed immediately for cell isolation. To dissociate cells from tumor and matched adjacent liver tissue, samples were perfused with digestion buffer containing 0.25mg/mL Collagenase IV (Sigma-Aldrich) and 0.1mg/mL DNase I (Sigma-Aldrich) in RPMI media with 10% FBS and finely minced. Tissues were enzymatically digested for 30min at 37°C with continuous agitation at 80rpm; The resulting cell suspension was mechanically disrupted using a 20mL syringe and 16G needle, then sequentially filtered through 100µm and 70µm cell strainers. Filters and tubes were rinsed with RPMI media with 10% FBS and, and cells were pelleted by centrifugation at 500g for 10min at 4°C. The cell pellets were resuspended in 25% Percoll (diluted in 10X PBS, Cytiva Sweden AB) layered over 70% Percoll, and centrifugated at 400g for 20min at room temperature (RT) with 0 acceleration and 0 break. The middle layer interphase was collected and washed with HBSS (Life Technologies). Red blood cells were lysed using 1X RBC lysis buffer (BioLegend) at RT for 2min, followed by an additional wash in HBSS. After a final centrifugation, cells were resuspended in buffer for counting and further profiling assays.

### Multiplex IHC imaging

Formalin fixed paraffin embedded tissues (FFPE) sections (4µm thick) were processed for multiplexed immunohistochemistry using the MICSSS (multiplexed immunohistochemical consecutive staining on a single slide) protocol, as previously described ^55^. In brief, slides were incubated overnight at 50°C to ensure proper adherence, then deparaffinized using xylene and rehydrated in decreasing concentration of ethanol (100%, 90%, 70%, 50%) followed by dH2O. Antigen retrieval was performed by heating slides in either pH 6 or pH 9 buffer at 95 °C for 30min. Slides were then treated with 3% hydrogen peroxide for 15min to quench endogenous peroxidase activity, and then incubated for 30min in a serum-free protein block solution (Dako). Primary antibody staining was performed using the optimized dilution, either for 1h at RT or overnight at 4°C, depending on the target. Signal amplification was achieved using horseradish peroxidase (HRP)-conjugated secondary antibody incubated during 30min, followed by chromogenic revelation using AEC substrate (Vector). Slides were counterstained with hematoxylin, mounted using a glycerol-based mounting medium, and scanned with an Aperio AT2 scanner (Leica). Following image acquisition, slide coverslips were removed by immersion in warm water (∼50°C) and the slides were bleached and subject to subsequent rounds of staining as previously described ^55^. Primary antibodies are presented in Table S2.

### scRNAseq hashing, processing and sequencing

Cell concentrations were obtained using the Nexcelom Cellaca. For each sample, 400,000 cells were aliquoted and centrifuged at 350g for 5min at 4°C. Supernatants were discarded, and pellets were resuspended in a solution containing hashtag-conjugated antibodies, prepared according to the New York Genome Center hashing protocol. Cells were incubated on ice for 20min for optimal binding. Unbound antibodies were removed through three sequential washes with 1mL of wash buffer (PBS + 0.5% BSA), and centrifugation at 350g for 5min at 4°C. Pellets were resuspended in 150µl wash buffer and counted again using the Nexcelom Cellaca. Hashed-labeled samples were then pooled, pelleted at 350g for 5min at 4°C, and resuspended in wash buffer to a final concentration of 4 x 10^6^ cells/mL. For scRNAseq (directly load), cells were loaded directly onto the 10x Genomics platform using either the 3’v2, 3’v3, or NextGem 5’v1 chemistry following the manufacturer’s instructions, with a targeted cell recovery of 10,000 cell per lane. Hashed sample pools were similarly loaded on the 10x Genomics NextGem 5’v1 assay, with a targeted cell recovery of 25,000 cells per lane. Gene expression and Feature Barcode libraries were prepared according to the 10x Genomics protocol. Hashtag oligonucleotides (HTO) were enriched during cDNA amplification by adding 3pmol of a HTO Additive primer (5’GTGACTGGAGTTCAGACGTGTGCTC). The PCR products were isolated from the mRNA-derived cDNA via SPRISelect size selection, and libraries were made as per the New York Genome Center Hashing protocol. All libraries were quantified using the Agilent 2100 hsDNA Bioanalyzer and KAPA library quantification kit (Roche). Sequencing of gene expression libraries was conducted on an Illumina NovaSeq 6000 platform, with a targeted depth of 25,000 reads per cell. HTO libraries were sequenced separately on an Illumina NextSeq 500 with a targeted read depth of 1,000 reads per cell. For 3’ libraries, paired-end sequencing was performed using 28bp for Read 1 (capturing UMI and cell barcode), 80 bp for Read 2 (transcript), and an 8bp i7 index read. For 5’ libraries, sequencing was done using 26bp for Read 1, 80bp for Read 2, and 8bp i7 index with 0-bp i5 read in either case.

### PICseq

Cells were resuspended in ice-cold sorting buffer (PBS supplemented with 0.2mM EDTA, 0.5% BSA) with with anti-human TruStain FcX (BioLegend) to block Fc receptors. Cells were then labeled with a panel of fluorochrome-conjugated antibodies targeting specific surface markers. The following antibodies were used: CD235a (Pacific Blue), TCRβ (FITC), CD11c (PE) and CD64 (PE, BioLegend); CD45 (APC, eBioscience) and CD19 (Pacific Blue) and CD56 (PE-Cy7, BD Biosciences). To assess cell viability, DAPI was added just prior to sorting. The different cell types of interest were sorted using an ARIA-III instrument (BD Biosciences), acquired using BD FACSDIVA software (BD Biosciences) and analyzed with FlowJo. Live cells were individually sorted into 384-well plates pre-loaded with 2μl of lysis buffer containing barcoded poly(T) reverse-transcription primers for scRNAseq ^56^. Each plate included four empty wells as negative controls for downstream data quality control. Immediately after sorting, each plate was centrifuged to ensure cells entered the lysis solution and were then stored at −80°C until further processing. ScRNAsea libraries were subsequently prepared using MARS-seq protocol, following the procedures previously described ^27,28^.

### Gene selection for MERFISH

To identify transcriptionally distinct cell population using MERFISH, we developed two sequential gene panels of informative transcripts. The gene selection strategy was divided into two main approaches. The first set consisted of manually curated genes serving as markers for various immune cell subsets including macs, T cells, B cells, and dendritic cells, as well as functional markers to capture biological processes such as T cell exhaustion, proliferation, signaling etc. The second set included genes identified as differentially expressed from prior scRNAseq datasets, selected to further resolve cellular heterogeneity. We evaluated candidate genes using the MERSCOPE Gene Panel Design Portal (Vizgen, portal.vizgen.com) to ensure optimal compatibility with MERFISH requirements. Specifically, we verified that each gene was of sufficient length to allow enough encoding probes to bind, and that the entire gene panel meets the abundance threshold to avoid optical crowding for MERSCOPE imaging. This ended in two final panels of 400 and 500 genes (Table S3). To serve as a control for unspecific binding of probes, we included 50 blank barcodes.

### Tissue preparation for MERFISH

#### Sectioning and permeabilization of fresh frozen tumor samples

HCC samples were snap frozen and preserved in optimal cutting temperature (OCT) compound and stored at −80°C until processing. Tissue sections were cut at 10µm thickness using a cryostat (Microm HM525, Thermo Scientific) maintained at −20°C and mounted onto MERSCOPE Slide (Vizgen 20400001). Sections were fixed with 4% paraformaldehyde in PBS for 15min at RT, washed three times with PBS, and then transferred to 70% ethanol at 4°C for overnight permeabilization.

#### Immunostaining and cell boundary labeling

Following permeabilization, sections underwent photobleaching for 4h using the MERSCOPE Photobleacher (Vizgen 1010003) to remove background fluorescence. Cell boundary visualization and immunostaining were performed using the Vizgen Cell Boundary Kit (10400009), according to the Vizgen User Guide for fresh and frozen tissue sample preparation (https://vizgen.com/resources/user-guides/). Briefly, tissue sections were blocked in Blocking Solution (PN 20300012) with RNase inhibitor (NEB, M0314L) at 1:20 dilution for one hour, washed with PBS and followed by sequential incubation with Cell Boundary Primary Antibody Mix (PN 20300010) with the supplement of RNase inhibitor (NEB, M0314L) at 1:20 dilution. After three washes with PBS, oligo-conjugated secondary antibodies were applied for 1h with RNase inhibitor (NEB, M0314L) at 1:20 dilution, followed by a 15min post-fixation in 4% paraformaldehyde. Sections were then washed with PBS and prepared for probe hybridization.

#### MERFISH encoding probe hybridization

Tissues were pre-washed with 5mL of Vizgen Sample Prep Wash Buffer (Vizgen 20300001) for 5min, then incubated in 5mL Formamide Wash Buffer (Vizgen 20300002) at 37°C for 30min. Hybridization was performed by applying 50µL of the custom MERSCOPE Gene Panel Mix (Vizgen 20300008, Table S3) directly onto the tissue, sealing with parafilm, and incubating at 37°C for 36-48 hours. Post-hybridization washes included two rounds with Formamide Wash Buffer at 47°C for 30min, followed by a 2min rinse with Sample Prep Wash Buffer.

#### Gel embedding and tissue clearing

Samples were embedded in a hydrogel composed of 5mL Gel Embedding Premix (Vizgen 20300004), 25µL of 10% ammonium persulfate (Sigma, 09913-100G) and 2.5µL of TEMED (N,N,N’,N’-tetramethylethylenediamine; Sigma, T7024-25ML). 20mm Gel Coverslips (Vizgen 20400003) were treated with RNaseZap, 70% ethanol and Gel Slick (VWR, 12001-812). After preincubating the tissue with gel solution for 1min, 100µL of fresh gel mix was applied, and samples were covered with a Gel Coverslip. After removing excess solution, samples were left at RT for 1.5 hours to polymerize. Gel Coverslips were then removed, and tissue sections were incubated in Clearing Solution (50µL of Protease K (NEB, P8107S) in 5mL of Clearing Premix (Vizgen 20300003)) first overnight at 47°C, followed by a second overnight incubation at 37°C. For complete details, refer to Vizgen’s online user guide for fresh and frozen tissues: https://vizgen.com/resources/fresh-and-fixed-frozen-tissue-sample-preparation.

#### Sample imaging

Following tissue clearing, samples were rinsed with Sample Prep Wash Buffer for 10min. To visualize nuclei and polyadenylated transcripts, sections were incubated with 3mL of DAPI and Poly-T Reagent (Vizgen 20300021) for 15min at RT. This was followed by a 10min wash with Formamide Wash Buffer and a final transfer to Sample Prep Wash Buffer. Imaging buffer was prepared by adding Imaging Buffer Activator (Vizgen 20300015) and RNase inhibitor to the Imaging Buffer Base. Processed samples and imaging reagents were loaded into the MERSCOPE system (Vizgen 10000001). A low-resolution DAPI mosaic was acquired at 10x magnification to select regions of interest, which were subsequently imaged at high resolution using a 60x objective. A full instrumentation protocol is available via Vizgen’s resource portal: https://vizgen.com/resources/merscope-instrument/. The resulting spatial transcriptomics data were used for cell segmentation and downstream analysis.

#### MERFISH in FFPE tumor samples

For FFPE samples, 5µm-thick sections from HCC tumors were cut using a microtome (Microm HM325, Thermofisher) and mounted onto on MERSCOPE FFPE Slides (Vizgen 20400100), following Vizgen’s MERSCOPE User Guide for FFPE samples (https://vizgen.com/resources/merscope-formalin-fixed-paraffin-embedded-tissue-sample-preparation-user-guide/). Sections were air-dried for 20min RT and the baked at 60°C for 10min. Deparaffinization was performed twice using Deparaffinization Buffer (Vizgen 20300112) at 55°C for 5min, followed by sequential ethanol washes (3x in 100% ethanol for 2min each, then 90% and 70% ethanol for 2min each). After rehydration, slides were incubated with Decrosslinking Buffer (Vizgen 20300115) at 90°C for 15min, cooled to RT, and treated with Conditioning Buffer (20300116) at 37°C for 30min. This was followed by a 2h incubation with Pre-Anchoring Reaction Buffer (20300113) at 37°C. Cell boundary staining was then performed using the Vizgen Cell Boundary Kit (10400009), following the same protocol used for fresh-frozen samples (https://vizgen.com/resources/user-guides/). Sections were washed with Formamide Wash Buffer (Vizgen 20300002) at 37°C for 30min and then incubated overnight at 37°C with Anchoring Buffer (Vizgen 20300117). Following this incubation, gel embedding and clearing steps were conducted as described for fresh-frozen tissues. Finally, samples were photobleached for 3h using the MERSCOPE Photobleacher (Vizgen 10100003), washed with Formamide Wash Buffer at 37°C for 30min, hybridized with the custom MERSCOPE Gene Panel Mix for 48h at 37°C, and imaged following the same workflow used for fresh frozen samples.

### Quantification and statistical analysis

#### Analysis of scRNAseq data

For scRNAseq libraries, raw sequencing data were processed using the Cell Ranger Single-Cell Software Suite (10X Genomics, v2.2.0), which performs sample demultiplexing, alignment, filtering, and UMI counting. Reads were aligned to the human GRCh38 genome assembly using the RefSeq gene model.

#### Batch-aware unsupervised clustering

Single cells isolated from tumor and adjacent tissue samples were filtered to retain cell barcodes with > 500 UMI, < 25% mitochondrial gene expression, and with less than defined thresholds of expression for genes associated with red blood cells and epithelial cells. Clustering was performed using an unsupervised batch-aware method previously described ^24^ with minor adjustments. This model, based on a multinomial mixture of gene expression distributions, incorporates a batch-specific noise term and distinguishes true cellular transcriptomes from sample-specific background. The clustering algorithm iteratively refines both cell-state assignments and noise estimates using a pseudo expectation-maximization (EM) approach ^57,58^. Unlike standard clustering methods, this framework enables robust integration of cells from multiple batches and allows classification of new datasets using the learned model. We applied this method to single cells derived from 73 tumor and 61 adjacent liver samples from 16 HCC patients, sequenced before February 28, 2021. Additional samples sequenced later were mapped onto the pre-trained model.

Model parameter estimation was performed using the pseudo-EM algorithm previously described ^59^, with the following modifications: (1) training and test set sizes were 4,000 and 2,000 cells, respectively; and (2) the optimal clustering seed was selected from 5,000 k-means+ runs. Genes contributing to patient-specific variability were excluded from the clustering analysis, including mitochondrial and stress-related genes, metallothioneins, immunoglobulin variable chains, HLA class I and II molecules, and three highly variable genes (MALAT1, JCHAIN, XIST). Ribosomal genes were excluded only during the k-means clustering step (step 2.D in Martin et al.) ^59^.

### Gene-module analysis

As previously described ^24,59^, cells were down-sampled to 2,000 UMI prior to variable gene selection. Gene-gene correlation matrices were computed and averaged across samples using Fisher Z-transformation. The resulting mean correlation matrix was then inverse-transformed to obtain consensus gene-gene correlation coefficients across the dataset. Genes were grouped into co-expression modules using complete linkage hierarchical clustering based on correlation distance.

### PICseq analysis

PIC deconvolution was performed as previously described ^27^, using the clustering model derived from the 10x Genomics scRNA-seq data as a reference. This enabled accurate mapping of PICs to our defined immune cell clusters. MNP abundance in PIC and singlet cells were quantified relative to the total population of DC and macs, excluding other myeloid subsets. Frequency comparisons between PIC-derived macs and their corresponding singlet mac populations were then performed separately for each cluster.

### MERFISH Cell Segmentation and Analysis

Tissue sections and probes were visualized using MERSCOPE Visualizer v2.1.2593.1. Cell segmentation was performed on nuclear and cytoplasmic stains using the Vizgen Postprocessing Tool (https://vizgen.github.io/vizgen-postprocessing/), with Cellpose v.1.0.2 applied across all available z-slices. The resulting 2D segmentations were integrated across slices to reconstruct 3D segmented cell. Consistent with previous reports ^12,60,61^, downstream clustering and analysis of small, transcriptionally similar, and spatially adjacent cell populations presented significant challenges. These issues stemmed both from biological similarities among interacting cell types and from segmentation inaccuracies, particularly for densely co-localized cells. Common analytical tools such as Seurat, Mutual Nearest Neighbors, Tangram, scANVI, and gimVI yielded variable performance across different cell types.

To improve label transfer from the annotated scRNA-seq atlas, Cellpose-derived segmentation masks were shrunk by 20% along both the x and y axes to remove transcripts located at the cell periphery, which are most prone to misassignment. While this improved transfer accuracy, it also introduced a systematic bias by eliminating peripherally localized transcripts, potentially reducing transcript diversity within cells. The shrunken segmentation masks were then used to partition transcripts via the Vizgen Postprocessing Tool. Single-cell expression matrices were analyzed using the Scanpy v1.9.1 and scvi-tools v1.0.4 Python libraries. Quality control filtering excluded cells with <10 or >750 transcript counts, <10 unique expressed genes, high blank probe signal (top 5th percentile), extreme spot densities (top and bottom 0.005 percentiles), and extreme Poly-T signal intensities (top and bottom 0.005 percentiles) to remove artifacts such as debris, apoptotic cells, and doublets. To integrate datasets across seven samples, an scVI model was trained using default parameters and incorporating tissue type (OCT vs FFPE), segmented cell volume, and patient ID as covariates. Cell type label transfer from the annotated scRNA-seq dataset was performed using TACCO (https://www.nature.com/articles/s41587-023-01657-3) with raw MERFISH counts as input. Assigned labels were cross-validated with Leiden clustering from the scVI-integrated data to ensure concordance. To spatially annotate tissue regions, cells identified as tumor cells, endothelial and hepatic stellate cells, and naïve or memory B cells were grouped into “tumor,” “stromal,” and “immune aggregate” compartments, respectively and drawn using Shapely v2.0.1 and GeoPandas v0.14.0. These regions were constructed using alpha shapes on centroid coordinates and buffered by 30 microns to account for sparse cell distribution. In cases of overlap, region priority was assigned in the following order: immune aggregate > stromal > tumor.

Co-occurrence analysis was performed separately for cells identified as TREM2 and FOLR2 mo-macs in tumor, stromal, and immune aggregate regions using squidpy v1.2.2 ^62^, using the squidpy.gr.co_occurrence() function at regular intervals from 0 to 500 microns and the TACCO assigned cell type labels as inputs. Enrichment values for each cell type at 30 microns were normalized for visualization by ln(1 + x), where x is the co-occurrence enrichment value as derived from squidpy. As a result of log transformation with pseudocounts, ln(2) (roughly 0.693) represents the threshold above which a given cell type is enriched, and below which a given cell type is de-enriched. Co-occurrence results were also checked for agreement with permutation-based neighborhood enrichment, calculated through the squidpy.gr.nhood_enrichment() function.

To identify additional gene expression programs using an orthogonal approach, consensus Non-Negative Matrix Factorization (cNMF) (https://elifesciences.org/articles/43803) was also carried out on the raw MERFISH counts derived from shrunken cell segmentation masks. cNMF v.1.4 was run with a range of k = 2 to k = 50 factors, with 100 independent solutions calculated per k, and 2,000 iterations per run. A k value of 33 was chosen for the best trade off of factor stability versus error. A local density threshold of 0.15 was selected to remove outlier runs, and remaining cNMF solutions for k = 29 were aggregated. Weights per gene and cell for each factor were calculated and visualized for analysis.

### Comparative analysis of external datasets

To determine transcriptional similitude of mac subsets across datasets, UMI counts were downloaded for each respective study (EGAS00001007547, GSE155698, GSE154826), matrices were normalized and scaled, and quality control metrics were applied; Barcodes matching cells with >500 UMIs were selected for and those with transcripts >25% mitochondrial genes were filtered from downstream analyses. Unsupervised clustering using a *K*-nn graph partitioning approach was performed and clusters of cells expressing mac genes were subsetted for follow-up analyses. Shared library genes were determined across studies, and matrices for select mac clusters were subsetted by the conserved library. Expression *z*-scores were computed within each dataset across the different mac clusters, based on mean UMI values per gene. Based on these *z*-scores, hierarchical clustering was applied to determine similitude across transcriptomes. Other R packages used include: scDissector, v.1.0.0; shiny, v.1.7; ShinyTree v.0.2.7; heatmaply v.1.3.0; plotly v.4.10.0; ggvis v.0.4.7; ggplot2 v.3.3.5; dplyr v.1.0.7; Matrix v.0.9.8; seriation, v.1.3.5.

### Quantification of MICSSS (MARQO)

For multiplexed imaging, we internally processed our samples to quantitatively analyze localization and coexpression patterns from MICSSS images. The tissue annotation was done by pathologists using QuPath software^63^. For the analysis, we used the svs multi-resolution, pyramidal images obtained per marker after staining. Each image underwent a brief quality control step to ensure tissue masking appropriately captured the tissue area. Both the AEC chromogen stain and the hematoxylin nuclear counterstain were extracted from each image via a dynamically determined deconvolution matrix. Then, each image was split into smaller tiles to permit computational analysis. Each tile from the first stained image was matched to the respective tile from the sequential stained images and then elastically registered using the extracted hematoxylin nuclear stain and SimpleElastix open-source software. Then, by using an iterative nuclear masking via STARDIST, we produced a composite semantic segmentation for nuclei residing in the series of tiles. Each nucleus was artificially expanded by a number of pixels to simulate a cytoplasm per cell and that was coherent with membrane marker staining. Lastly, cellular-resolution metadata was acquired for all cells in the final cell mask, including AEC and hematoxylin intensity properties (percentiles, dynamic ranges, etc.) and morphological characteristics (circularity, area, etc.) per cell. A final data frame was appended for each tile processed, producing one data frame representing all the cellular metadata per sample. To unbiasedly determine positive and negative cells per marker, we used an unsupervised classification technique to cluster cell populations, followed by a supervised approach where we would evaluate each cluster as positive or negative per marker. First, the metadata aggregated per sample was collected, transformed to z-scores, randomized, and split into subsamples per batch. Each batch was processed in parallel: data for each marker was transformed, clustered and collapsed into multiple groups by principal components analysis and uniform manifold approximation and projection. Then, we performed the final quality control to manually attribute which clusters were positive or negative. This produced a final cellular-resolution data frame containing binary marker classification that was used for downstream localization, marker coexpression, tissue annotation and reconciliation and statistical analyses.

### Statistical analysis

Statistical analyses were performed using Prism V.10 (GraphPad, California, USA). Survival data were analyzed using the Kaplan-Meier and log-rank tests for survival distribution. Multigroup analyses of variances were performed using one-way or two-way analysis of variance (ANOVA) followed by indicated post-tests. For simple comparison analysis, the unpaired Student’s t-test was used to compare parametric distributions.

## DATA AVAILABILITY

Human sequencing and MERFISH data will be available without restrictions at the time of publication.

## ACKNOWLEDGEMENTS

This study was funded by Regeneron Inc. We thank members of the Merad and Brown laboratories at the Marc and Jennifer Lipschultz Precision Immunology Institute at Mount Sinai and the Tisch Cancer Institute for insightful discussions and feedback; and the Mount Sinai Flow Cytometry Core, the Human Immune Monitoring Center (HIMC), the Center for Comparative Medicine and Surgery for animal husbandry and Biorepository and Pathology CoRE Laboratory at the Icahn School of Medicine at Mount Sinai for support. We thank the patients and their families for participating in the clinical trials. This work was supported in part through the computational resources and staff expertise provided by Scientific Computing at the Icahn School of Medicine at Mount Sinai. P.H. received the SITC-NanoString Technologies Spatial Profiling Award for the GeoMx assays. N.M. was partially supported by Fondation pour la Recherche Medicale (FRMSPE201803005095) and College des Enseignants en Dermatologie de France. S.G. was partially supported by National Institutes of Health (NIH) grants CA224319, DK124165, CA263705 and CA196521. A.O.K. and T.U.M. were supported in part by the Tisch Cancer Institute Cancer Center Support Grant (P30 CA196521). A.O.K. and E.H. were supported in part by R01 AI153363. M.M. was partially supported by NIH grants CA257195, CA254104 and CA154947. R.M. was supported by the 2021 AACR-AstraZeneca Immuno-oncology Research Fellowship, Grant Number 21-40-12-MATT. A.L. was supported by NIH/NCI R37CA230636, NIH/NCI R01CA251155, and Damon Runyon-Rachleff Innovator Award. S.W. was partially supported by a grant from Boerhinger Ingelheim. Figure 1A was Created with BioRender.com.

## AUTHOR CONTRIBUTION

P.H., G.T., M.S., T.U.M. and M.M. conceptualized the project. M.M. and T.U.M. obtained funding for the project. P.H. and M.M. designed the experiments. P.H., M.D.P., J.LB., M.B., K.L., A.T., R.Mattiuz, T.D., L.T., G.I., C.P., M.C., O.B., A.Reid, C.C., R.Marvin, H.S., G.C., R.Merand, L.M.G, M.N., Y.W., N.M., F.D. and Y.L. performed experiments. P.H., M.D.P., J.LB., A.M., B.Y.S., F.R., M.C., A.G. and S.C. analyzed experiments. M.S. and T.U.M. provided clinical care to patients in the clinical trial. Z.Z., S.O., S.C.W. and MI.F. provided expertise for pathological response assessment and tissue annotation. C.H., A.L. and L.W. coordinated the clinical and research teams, and managed clinical specimens with the help of H.J., N.Y., T.C., S.WI., H.B., and N.J. M.D.P., A.M., B.Y.S., M.B., I.F., D.D., F.R., A.G., S.C., S.H., J.K. and V.R. performed computational molecular and spatial analyses. R.Mattiuz, E.G-K., N.F., S.H., G.A., Y.L., A.Rahman, J.D., D.L., E.K., A.O.K., J.H., I.A., S.K-S., S.G., J.C.L., A.L., provided intellectual input. P.H. wrote the manuscript. M.D.P. and M.M. edited the manuscript. All authors provided feedback on the manuscript draft.

## SUPPLEMENTARY TABLE LEGENDS

Supplementary Table 1: Clinical and demographic data of patient cohorts

Supplementary Table 2: Primary antibodies for MICSSS

Supplementary Table 3: MERSCOPE Gene Panels

**Figure S1:**
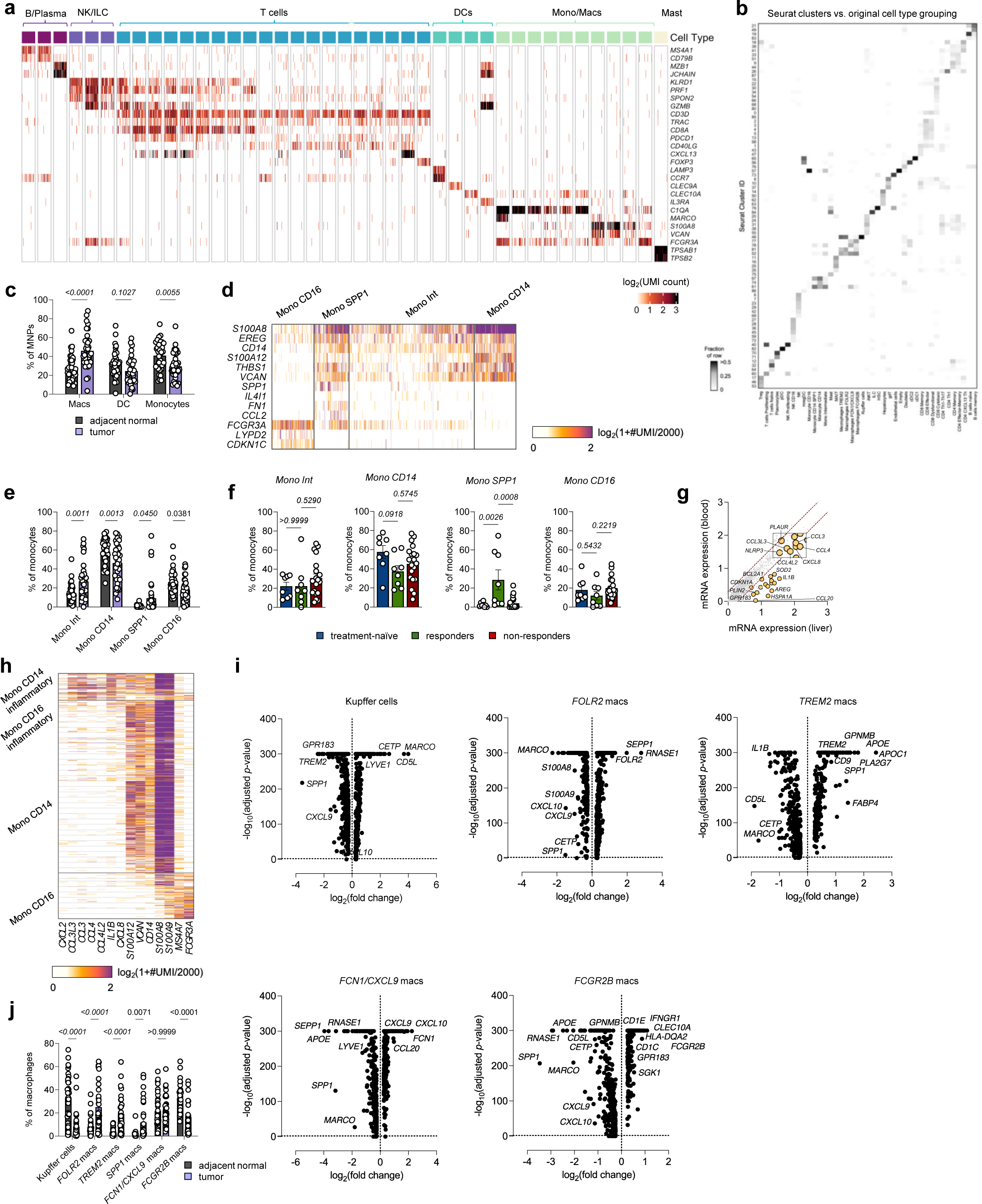
High resolution characterization of mononuclear phagocytes in early-stage HCC (related to figure 1) **a,** Expression of genes discriminating scRNAseq immune clusters showing number of UMI per cell. **b,** Mapping of cell type annotations (row) to Seurat clusters (column), with shading indicating fraction of row. **c,** Percentage of mononuclear phagocytes (MNPs) in adjacent and tumor tissues. Dots represent mean ± SEM. Two-way ANOVA followed by Sidack multiple comparison test was performed for each subset. **d,** Heatmap representing single-cell gene expression in monocyte clusters from both tumor and adjacent tissues. **e,** Percentage of monocytes in adjacent and tumor tissues. Dots represent percentage of monocytes ± SEM. Two-way ANOVA followed by Sidack multiple comparison test was performed to compared monocyte abundance in adjacent and tumor tissue for each subset. **f,** Percentage of monocytes in treatment-naïve, responders and non-responders to PD-1 blockade for each subset. Dots represent mean ± SEM. One-way ANOVA followed by Dunnett’s multiple comparison test was performed for each subset. **g,** Comparative analysis of gene expression of classical monocytes in tissue (adjacent and tumor) versus blood (PBMC). **h,** Heatmap representing single-cell gene expression of monocyte from circulating blood (PBMC). **i,** Volcano plots representing differential gene expression for each mac subset compared to all macs. **j,** percentage of macs in tumor and adjacent tissue for each subset of macs. Dots represent percentage of macrophages ± SEM. Two-way ANOVA followed by Sidack multiple comparison was performed.

**Figure S2:**
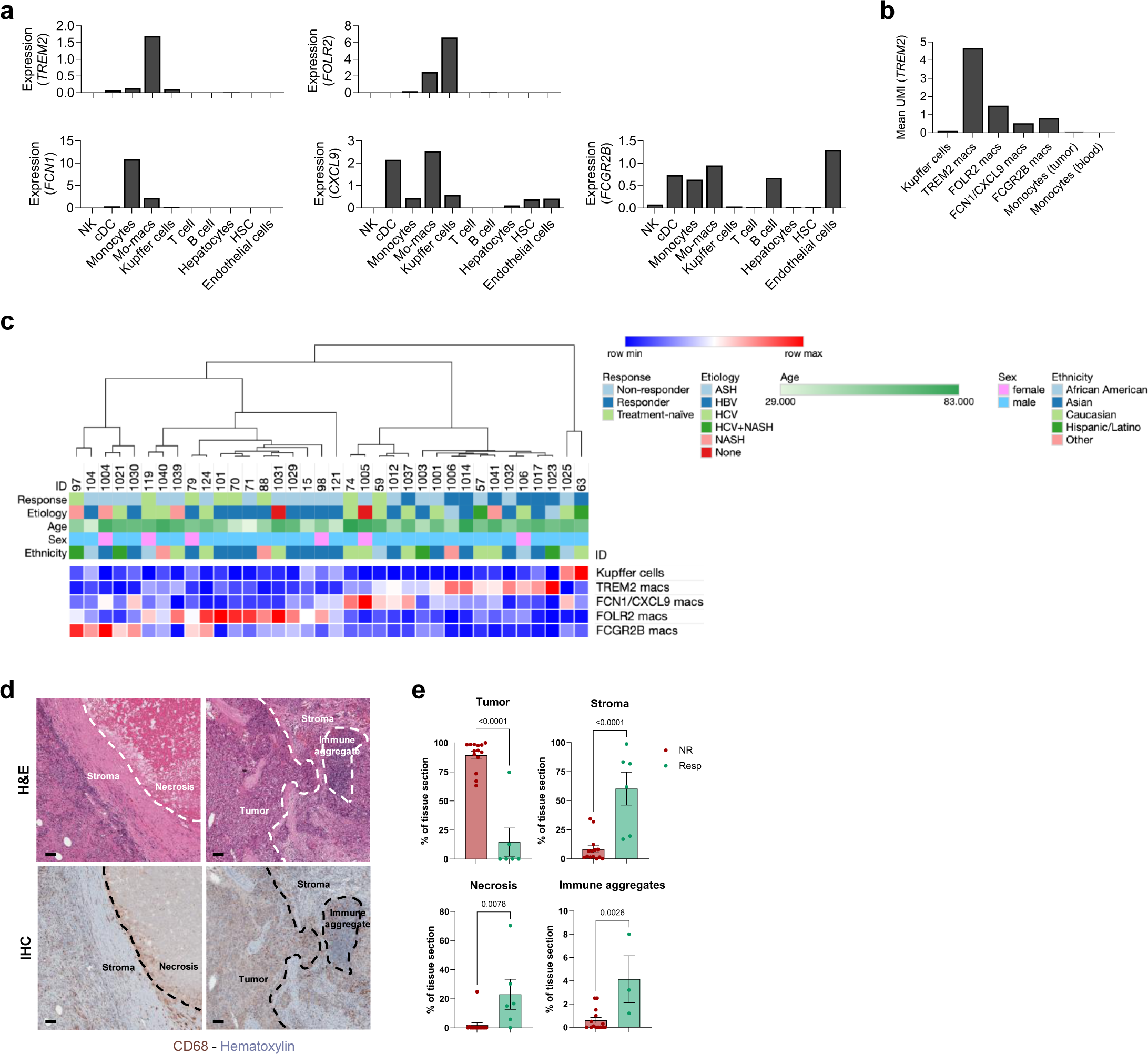
Monocyte-derived macrophage programs are specifically enriched in early-stage HCC (related to figures 1 and 2) **a,** Mean UMI expression of selected mo-macs transcripts *TREM2*, *FOLR2* (top) *and FCN1, CXCL9, FCGR2B* (bottom) across immune and non-immune populations. **b,** Mean UMI of TREM2 expression in the different subsets of macs and monocytes. c, Heatmap associating mac subsets with patients and associated clinical metadata showing that the main driver patient classification is treatment response. **d,** Representative images of the different ROI regions (immune aggregate, stroma, tumor nodule) selected for spatial transcriptomic using immune markers (CD3, CD68), stroma marker (aSMA) and nuclei staining (DAPI). **e,** Quantification of the different tissue area in responders and non-responders, including tumor nodules, stroma, necrosis and immune aggregates. Dots represent mean ± SEM. Unpaired t-test was performed for each graph.

**Figure S3:**
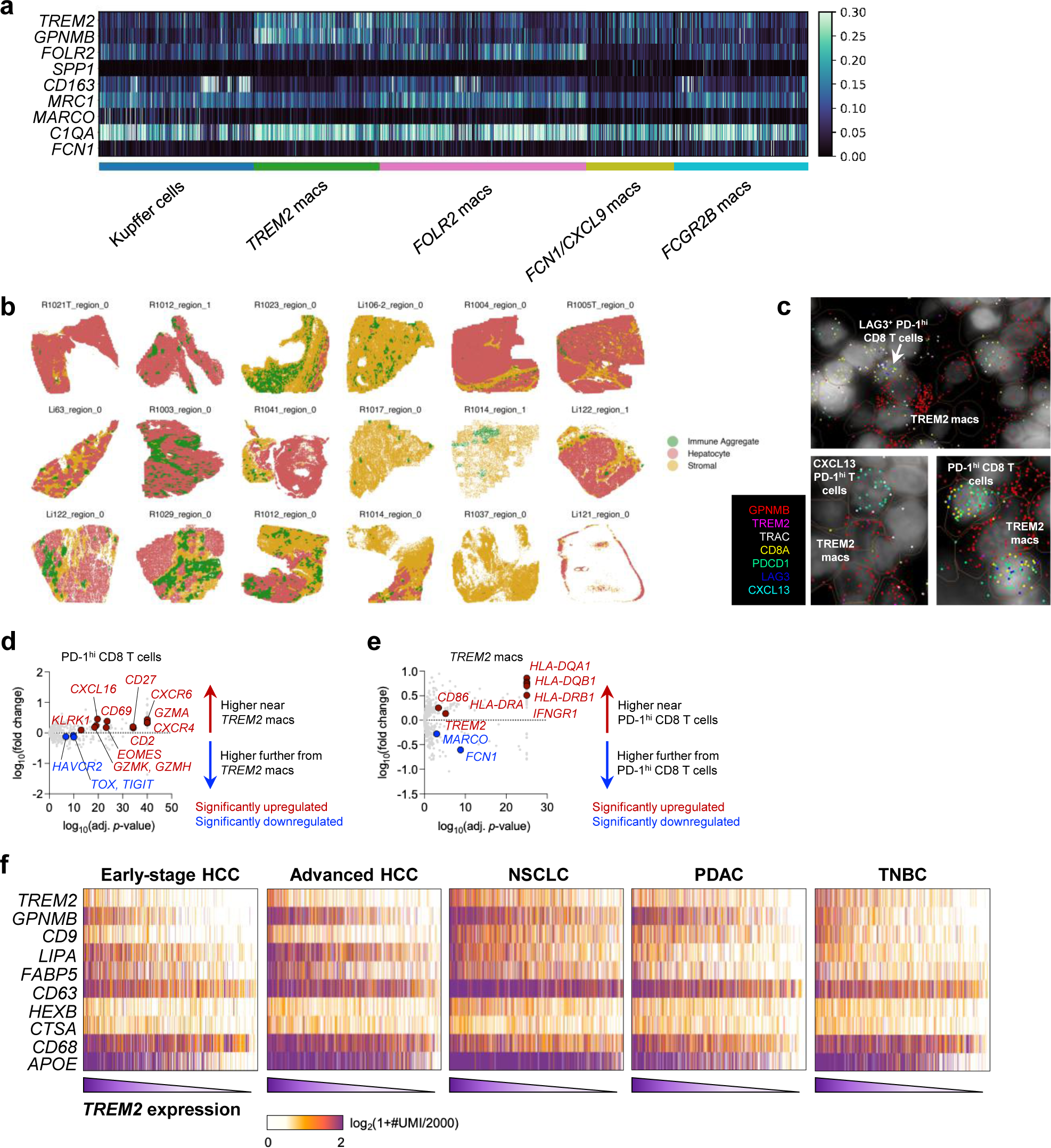
Spatial transcriptomics reveal intercellular interactions involving TREM2 macs (related to figures 2 and 3) **a,** Thruthplots representing gene expression on segmented clustered cells from MERFISH HCC samples. **b,** Tissue annotation of each HCC MERFISH samples. **c,** Representative images of TREM2 macs colocalized with different subsets of PD1hi T cells in tumor samples. Each dot represents a transcript. Brown circles represent cells from segmentation. **d,** Differential gene expression (DEG) between PD-1^Hi^ CD8 T cells in close proximity of TREM2 macs or further away from TREM2 macs. In red and blue are highlighted genes that were significantly upregulated or downregulated, respectively. **e,** DEG between TREM2 macs in cells in close proximity of PD-1^Hi^ CD8 T cells or further away from PD-1^Hi^ CD8 T cells. In red and blue are highlighted genes that were significantly upregulated or downregulated, respectively. **f,** Truthplot of *TREM2* program expression by all mo-macs across the different tumor types and ordered by *TREM2* expression.

**Figure S4:**
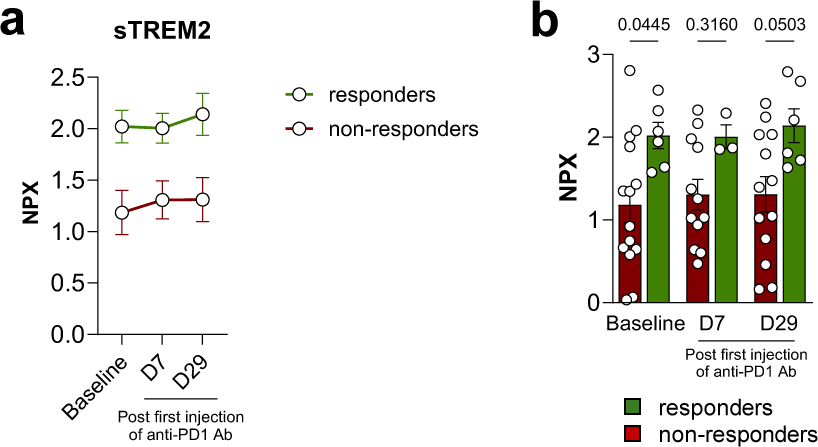
Soluble TREM2 is a biomarker of immunotherapy response (related to figure 4) **a-b,** Kinetic of normalized protein expression (NPX) of soluble TREM2 prior and during anti-PD-1 therapy in responders and non-responders grouped **(a)** and per sample **(b)**.

